# Disrupting Splicing Regulation to Rescue β-Catenin: A Novel Approach for Treating CTNNB1-Haploinsufficiency Disorder

**DOI:** 10.1101/2025.04.01.646581

**Authors:** Matea Maruna, Petra Sušjan-Leite, Maja Meško, Roman Jerala

**Author notes:** Corresponding author (RJ).

## Abstract

Loss-of-function mutations in the *CTNNB1* gene cause β-catenin deficiency, resulting in CTNNB1 syndrome—a rare neurodevelopmental disorder characterized by motor and cognitive impairments. Given the wide variety of mutations across *CTNNB1* and its dosage sensitivity, a mutation-independent therapeutic approach that preserves endogenous gene regulation is critically needed. This study introduces spliceosome-mediated RNA trans-splicing as a novel approach to restore β-catenin production. Precursor mRNA trans-splicing molecules (PTMs) targeting CTNNB1 introns 2, 5, and 6 were designed and evaluated using a split fluorescent YFP reporter system. Rationally designed short antisense RNAs, which mask splicing regulatory elements, significantly enhanced PTM-mediated trans-splicing at both RNA and protein levels. Additionally, introducing a self-cleaving ribozyme at the PTM’s 5’ end further improved trans-splicing efficiency, likely due to increased nuclear retention. CMV promoter-driven PTM expression yielded the highest efficiency. Importantly, successful trans-splicing of the endogenous CTNNB1 transcript confirmed the physiological relevance of this strategy. This study is the first to apply and optimize SMaRT for *CTNNB1* correction, providing a promising, mutation-agnostic approach for treating CTNNB1 syndrome.

## Introduction

Loss-of-function mutations in the catenin beta 1 (CTNNB1) gene lead to β-catenin deficiency, which is strongly associated with the CTNNB1 syndrome, a rare, monogenic, neurodevelopmental disorder marked by an array of motor and cognitive impairments with an estimated prevalence of around 2.6-3.2 in 100,000 live births^1^. Due to the haplo-insufficiency of the CTNNB1 gene, the pathological phenotype is driven by a single-allele mutation^2–4^. CTNNB1 mutations vary in type and position and can be found scattered across the whole coding and noncoding region of the CTNNB1 gene. This advocates the need for a generally applicable approach, capable of correcting large fraction of diverse mutations, including point mutations and indels ^4–6^. Furthermore, given the CTNNB1 pathogenicity in both deficiency, which leads to CTNNB1 syndrome and overexpression, which may lead to cancer, CTNNB1 gene bears characteristics of a dosage-sensitive gene, which suggests that its endogenous gene regulation should preferably be maintained^7–9^. Finally, it is currently unknown whether the mutated CTNNB1 transcripts undergo nonsense-mediated RNA decay or translate into truncated variants with potential dominant-negative effects that could hinder the function of the WT β-catenin from the remaining wild-type allele. Therefore, in some patients, gene upregulation and elimination of the mutated transcript might be necessary to fully restore CTNNB1 function.

One of the major processes of mRNA maturation is RNA splicing in which exons within the nascent RNA are ligated following intron removal. Splicing is catalyzed by the spliceosome, a large ribonucleoprotein complex, whose formation is orchestrated by five small nuclear RNA (U1, U2, U4, U5 and U6 snRNA) acting in concert with small nuclear ribonucleoprotein particles (snRNPs) and other protein factors^10^. The majority of pre-mRNA splicing proceeds in *cis* manner, with exon ligation occurring within the same pre-mRNA molecule. It can, however, also proceed in *trans* manner, where exons of different pre-mRNA molecules are combined into a hybrid mRNA in a process known as trans-splicing^10^. Naturally occurring trans-splicing which has been observed in trypanosomes^11,12^ and nematodes^13^ has inspired the development of an artificial exon replacement strategy – spliceosome-mediated RNA trans-splicing (SMaRT) with prospects as a research and therapeutic tool for the RNA repair^14^.

SMaRT is based on the rational design of an exogenous pre-trans RNA molecule (PTM) capable of replacing selected exons in a target pre-mRNA molecule, thereby generating a chimeric mRNA (Figure 1A). Depending on the region to be replaced in the target pre-mRNA, there are three types of trans-splicing: 3’, 5’ and internal trans-splicing^15^. SMaRT strategy allows for a partial gene replacement, which is particularly beneficial in the correction of large genes unsuitable for introduction through AAV vectors, correction of genetic defects regardless of their size and type and, most importantly, it maintains the endogenous transcriptional regulation of the target gene in terms of time, space and transcriptional output. To date, trans-splicing has been shown in several cell models of dominant and recessive genetic diseases such as cystic fibrosis^16,17^, spinal muscular atrophy^18^, Duchenne muscular dystrophy^19,20^, dystrophic epidermolysis bullosa^21,22^, and retinitis pigmentosa^23,24^. Furthermore, trans-splicing was recently reported as an elegant tool to facilitate the reconstitution of a split gene following dual AAV vector delivery^25^. While these successes highlight trans-splicing as a promising therapeutic strategy, its dissemination, both clinically and as a biochemical research tool, has been significantly hindered by its low efficiency ^26^. Several optimizations to the original PMT design, such as the selection of a suitable target intron and binding domain, the position and length of the binding domain, the presence of mismatch mutations, the thermodynamic stability of PMT terminal ends, and most recently, implementations of the CRISPR/Cas13 system, can significantly improve trans-splicing efficiency^27–31^.

**Figure 1.**
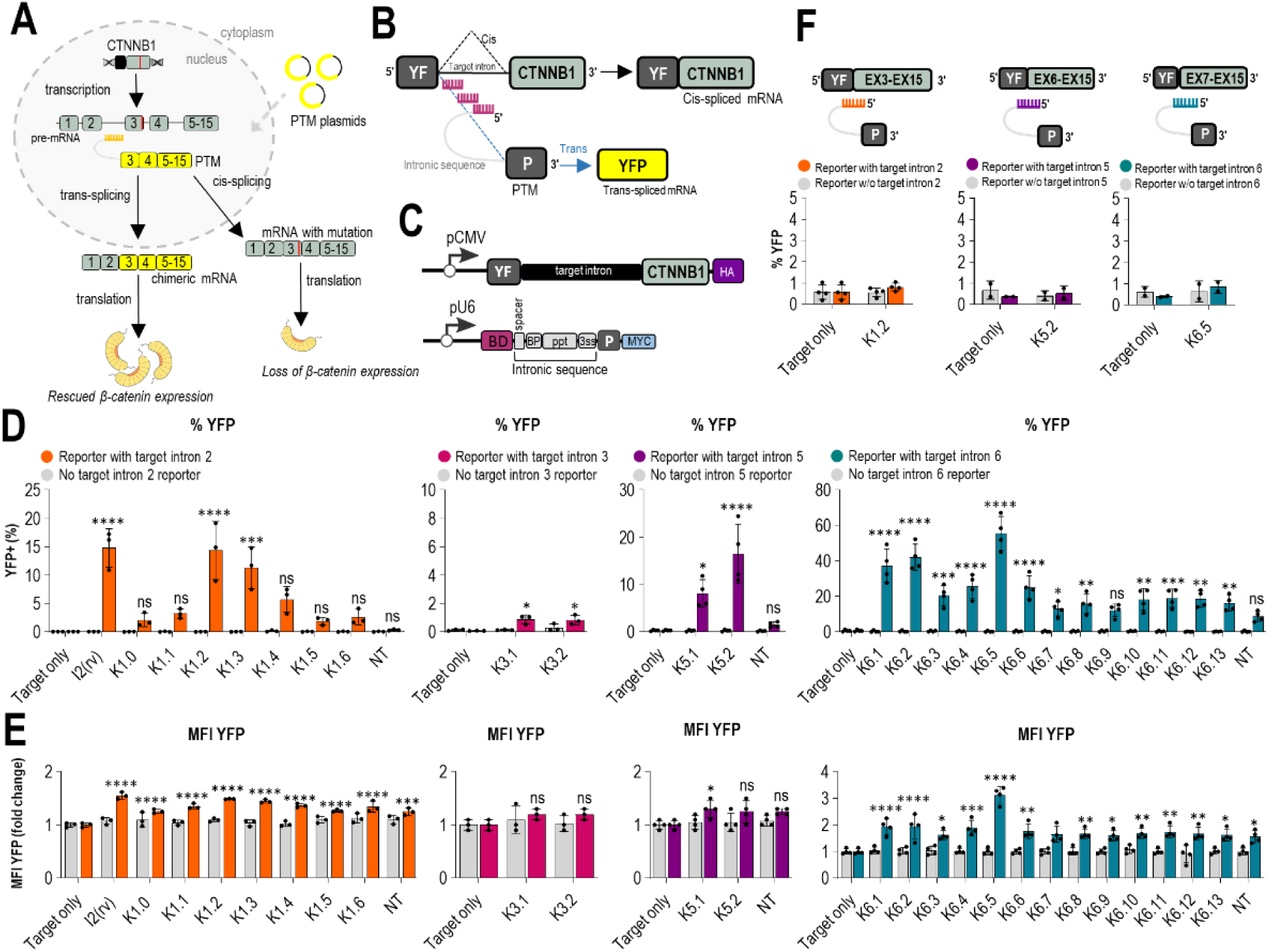
Trans-splicing mechanism and PTM candidate screening using a split YFP target intron reporter for efficient trans-splicing. (A) Following transcription, the CTNNB1 gene is transcribed into pre-mRNA containing a CTNNB1-associated mutation, leading to the loss of β-catenin expression. A pre-trans-splicing molecule (PTM) is introduced into the cells to induce trans-splicing and outcompete cis-splicing, resulting in the formation of chimeric mRNA without the mutation, thereby rescuing β-catenin expression. (B, C) Composition of the split YFP target intron reporter: It consists of the target mRNA and PTM candidate. The target mRNA, expressed under the CMV promoter, contains the N-terminal domain of YFP, a target intron, and downstream exons. The PTM molecule, expressed under the U6 promoter, includes a binding domain (BD), intronic sequence (spacer, branch point (BP), polypyrimidine tract (ppt), and 3’ splice site (3’ss)), the C-terminal domain of YFP, and a Myc tag. Accurate trans-splicing results in the fusion of both YFP domains into a functional protein, detected as fluorescence via flow cytometry. (D, E) Quantitation of flow cytometry for PTM screening targeting introns 2, 3, 5, and 6 was performed using the split YFP target intron reporter. Negative controls included PTM NT with a randomly generated binding domain, target intron reporter transfected alone, and PTM candidate transfected without the target intron reporter. Quantitation by flow cytometry was presented as the percentage of YFP+ cells and the median fluorescence intensity (MFI) normalized to the target intron reporter only control for the four different introns—2, 3, 5, and 6. Data are presented as the mean value ± SD from at least three independent experiments. Comparison to target intron reporter only (target only) was analyzed using one-way ANOVA with Dunnett’s multiple comparison test. ****p < 0.0001; ***p < 0.001; **p < 0.01; *p < 0.05; non significant (ns).

This study is the first to investigate the feasibility of SMaRT strategy for the replacement of CTNNB1 exons. Several CTNNB1 characteristics highlight trans-splicing as a promising avenue to pursue in search of treatment for the CTNNB1 syndrome. The above-mentioned issues regarding CTNNB1 could be elegantly avoided with a trans-splicing-based intervention, as it would eliminate the effect of diverse mutations and restore functionality through the exon replacement while retaining the endogenous gene regulation. We introduced both the initially reported PMT design and additional modifications aimed to enhance SMaRT efficiency. Upon PTM screening within a split YFP reporter system, we identified the best PMT candidates for three target introns within the CTNNB1 gene, with trans-splicing efficiency ranging from 15 % to 55 %. Trans-splicing efficiency was strongly enhanced by the addition of the rationally designed short antisense RNAs, expressed in a hybrid U1-U7 snRNA cassette that target splicing regulatory elements and inhibit cis-splicing. Furthermore, the choice of the CMV promotor for the PMT expression and novel modification of positioning a self-cleaving ribozyme on its 5’ significantly improved PMT-mediated trans-splicing targeting introns 2 and 5. This improvement allowed us to achieve the trans-splicing efficiency of the Cas13 by implementing trans-splicing techniques in the reporter system. For these introns, we were also able to detect robust trans-splicing of the endogenous CTNNB1 transcript indicating the physiological and potentially therapeutic relevance of our findings.

## Results

### Design of the PMT candidates supporting trans-splicing of CTNNB1 introns

In this study, we wanted to optimize 3’ trans-splicing for CTNNB1 gene that would maintain the endogenous transcriptional regulation and could correct the majority of reported mutations that cause CTNNB1 syndrome (Figure 1A). As the introns with stronger splice sites are more likely to favor cis-splicing and outcompete PMT^26^, we first identified the most suitable CTNNB1 intron candidates for trans-splicing based on the computational prediction of their splice site strength using MaxEnt program^33^. We selected CTNNB1 introns 2, 3, 5, and 6 as trans-splicing targets as they exhibited low MaxEnt scores, while strong 3’ splice sites were introduced in the designed PTM candidates (Table S1). Furthermore, targeting intron 2 would allow replacement of the majority of the coding region of the CTNNB1 transcript, making this approach universally applicable to all CTNNB1 syndrome-associated mutations, which are spread across the entire gene. Similarly, the effective targeting of introns 6 and 5 could be applied for many of the reported mutations, as the majority of these mutations are located in exons 7 and 8^4^. Previous studies have shown that the selection of suitable binding domain (BD) within an intron preceding the exons to be replaced is crucial in the design of an efficient PTM^31,34^. To this end, we utilized a tiling screening in which we designed PTM candidates with 150-nucleotide-long BDs spanning the entire sequence of target introns 2, 3 and 6 with 50-nucleotide overlaps. In terms of targeting intron 2, we also designed a binding domain as a reverse complement of the entire intron 2. As intron 5 is shorter, the BD length for intron 5 screening was 50 nucleotides (Figure 1B), with 15 nt overlap. Eight PTM molecules differing in their BD, complementary to different sites within the target intron, were screened for intron 2, fourteen for intron 6, and two PTMs for introns 3 and 5 (Figure S1).

To screen for PTM candidates with the highest trans-splicing efficiency, we designed a split (YFP) reporter system which consisted of the construct comprising target CTNNB1 intron (hence target intron reporter) and PMT in fusion with N- and C-terminal parts of the YFP gene, respectively (Figure 1B). Target intron reporter comprised the N-terminal domain of the YFP gene, target intron, downstream exons from the CTNNB1 gene, and the hemagglutinin (HA) tag (Figure 1C). PTM candidates were designed for 3’ trans-splicing and comprised BD, spacer, an intronic sequence including branch point, polypyrimidine tract (ppt), and 3’ acceptor splice site (3’ ss), followed by the C-terminal domain of the YFP gene and myc tag (Figure 1C). In this reporter system, trans-splicing between the target intron reporter and PTM molecule in cells is expected to assemble a functional YFP whose fluorescence can be measured by flow cytometry (Figure 1B).

Upon co-transfection of the target intron reporter and corresponding PTM into HEK293T cells in molar ratio 1:2, flow cytometry was used to measure the percentage of cells expressing YFP and median fluorescent intensity (MFI) of the YFP fluorescence. Specifically, cells were gated for the 20,000 iRFP+ cells, which served as a control for transfection efficiency, and were then further analyzed for YFP expression, indicating successful trans-splicing (Figure S6). An increase in the percentage of YFP^+^ cells from 15% to over 40% was detected with the best PMT candidates (K1.2, K6.5 and K5.2) in comparison to the control (Figure 1D). Accordingly, we also detected an increase in the YFP MFI, especially in the case of intron 6, where we detected a 3-fold increase (Figure 1E). Neither the target intron reporter, nor PTM showed any increase in YFP fluorescence when transfected alone (Figure 1D and 1E). Importantly, the signal was tightly associated with the presence of the BD with base complementarity to the target intron as a random RNA sequence in place of the BD resulting in little to no increase in the YFP signal compared to the tested PTMs (Figure 1D and 1E). Furthermore, we also performed co-transfection of the best PMT candidates (K1.2, K6.5 and K5.2) with a reporter lacking target intron to control for any spontaneous association of the translated N- and C-terminal YFP as a result of the overexpression. No increase in YFP signal was observed, indicating the absence of interaction between translated N- and C-terminal YFP in HEK293T cells (Figure 1F). PMT candidates targeting intron 3 did not lead to an increase in the percentage of YFP^+^ cells or MFI and were excluded from further experiments (Figure 1D and E).

We also investigated the effects of the length of the BD. Initially, we designed 150 nt BDs, which were reported before to provide the best trans-splicing efficiency, and least off-target effects in comparison to shorter or longer BDs^31^. Stability of substantially shorter RNA duplexes, such as 100 or even 50 nt are expected to be similar to the 150 nt. Therefore, to assess the impact of shorter BD, we shortened the BDs of best-performing PTM candidates, K1.2 and K6.5 to 100 nt and 50 nt. We tested truncations from the 5’, 3’ and both ends of the BD. However, all BD truncations led to decrease of the trans-splicing efficiency in all tested PTM candidates (Figure S2C and S2D). An explanation could be that longer BDs may be able to disrupt stable secondary structures within the mRNA through strand displacement where longer BD have higher probability to anneal to the single stranded regions within the mRNA than shorter segments.

### Antisense RNA strongly enhances PMT-mediated trans-splicing

We next aimed to improve the trans-splicing efficiency by introducing various modifications to the original PMT design to either inhibit the cis-splicing, enhance its nuclear retention, prevent PTM degradation or prevent translation of PMT in the absence of splicing. Combination of PMT with randomly generated antisense sequences and synthetic antisense oligonucleotides targeting specific intron-exon junctions was reported to boost trans-splicing, presumably by inhibiting the competitive cis-splicing^22^. To suppress cis-splicing, we opted for a design of short antisense RNAs (asRNA) with complementarity to the segments that contain BP and exon splicing enhancers (ESE) within the target intron and the first downstream exon (Figure 2A). To support robust asRNA expression combined with enhanced nuclear retention and resistance to degradation, asRNAs were expressed in a hybrid U1-U7 expression cassette in which the 5’ end of the modified U7 snRNA was replaced by the asRNAs of choice and expressed under a strong, DNA polymerase II-dependent snRNA U1 promotor^32^ (Figure 2B). We designed 30-nucleotide long asRNAs masking BP and 3’ splice site in the target intron-exon junction and 30-35-nucleotide long asRNAs masking selected ESE within the first downstream exon (Figure 2C). Designed asRNA were co-transfected to HEK293T cells in combination with the best corresponding PMT from screening in Figure 1D and 1E and the target intron reporter. Flow cytometry revealed that the addition of short asRNA significantly enhanced trans-splicing efficiency of PMT candidates for all target introns as observed both in percentage of YFP^+^ cells and MFI (Figure 2D and 2E). The asRNA candidate BP4 in the case of intron 2 trans-splicing increased the percentage of YFP^+^ cells from 19% to 48%, while the best asRNA for intron 6 trans-splicing (SF2/ASF) improved the YFP^+^ cells from 44% to 65%. In case of intron 5 trans-splicing, asRNA SC35 targeting exon 6 increased the YFP^+^ cells from 16% to 55% (Figure 2D). Improved YFP production was also observed in terms of MFI, with the highest increase detected for intron 6 (Figure 2E). We next wanted to see if the expression of multiple asRNA would further enhance PMT-mediated trans-splicing in any of the target introns. However, a combination of two or three asRNAs, expressed from a single vector, leads to no further improvement in trans-splicing efficiency in the split YFP reporter system (Figure S2A and S2B).

**Figure 2:**
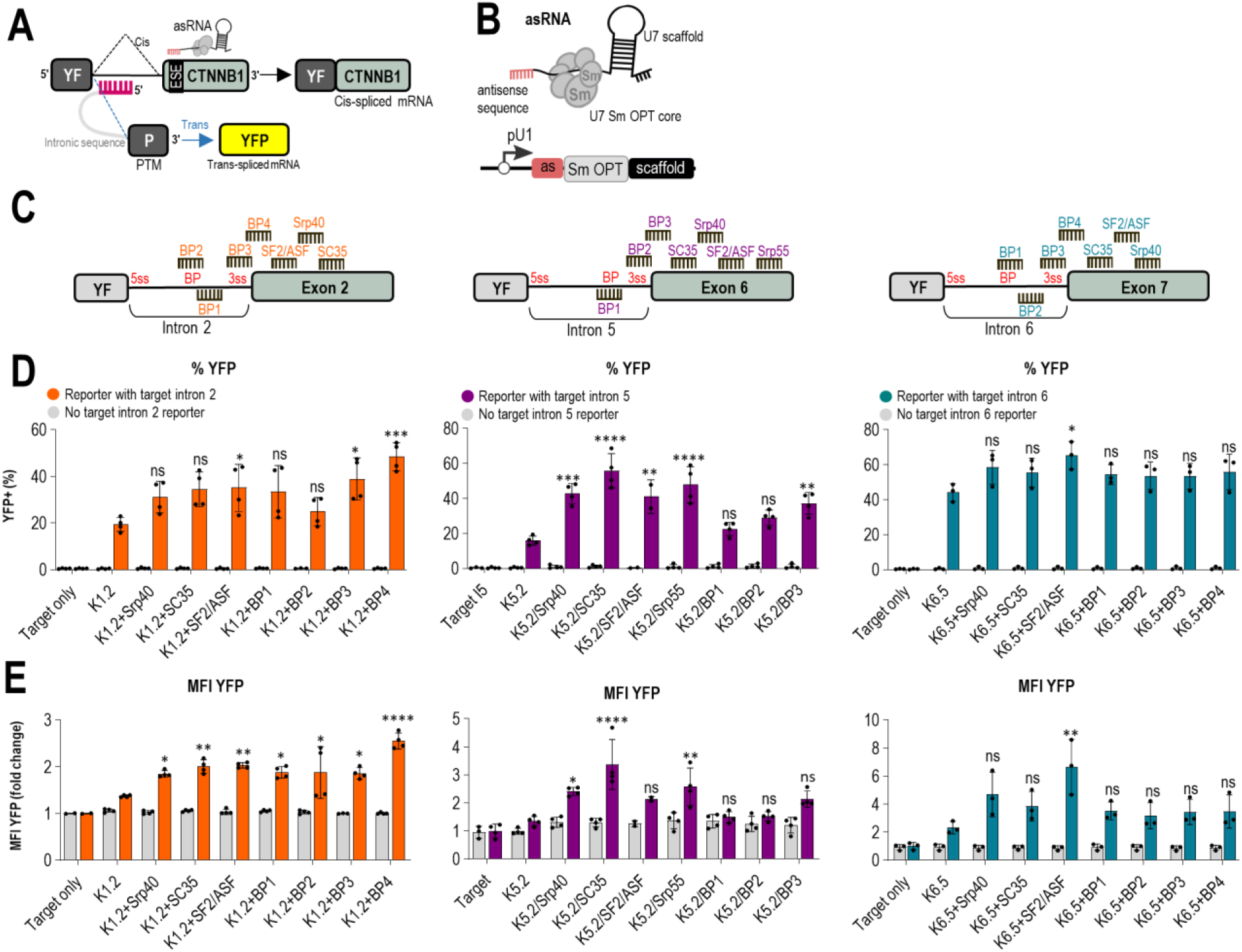
Short antisense RNAs enhance trans-splicing efficiency for CTNNB1 introns in target intron reporter. (A) A schematic representation illustrates the mechanism by which short antisense RNAs (asRNAs) block cis-splicing. The 30-35 nucleotide long asRNAs are designed to target regions from the branch point (BP) to the 3’ splice site in the target intron, as well as exonic splicing enhancers (ESEs) within the first exon downstream of the target intron. (B) The short asRNAs consist of an antisense sequence coupled with a modified U7 Sm OPT core, which binds Sm proteins found in spliceosomal snRNAs, along with a U7 snRNA scaffold. These chimeric U7-antisense sequences are expressed under the control of a strong pol III U1 snRNA gene promoter and termination sequence. (C) The positions of short asRNAs targeting splice sites and ESEs are mapped in target introns 2, 5, and 6, and their respective first downstream exons 3, 6, and 7 (D, E). The detection of enhanced trans-splicing efficiency by adding short asRNAs was performed using flow cytometry. Negative controls included the target intron reporter transfected alone, and the best PTM candidates (K1.2, K5.2, and K6.6) co-transfected with asRNA but without the target intron reporter. Quantitation by flow cytometry is presented as the percentage of YFP+ cells (D) and the median fluorescence intensity MFI (E) normalized to the reporter-only control for the three different introns—2, 5, and 6. Data are presented as the mean value ± SD from at least three independent experiments. Comparison to PTM candidate co-transfected with target intron reporter was analyzed using one-way ANOVA with Dunnett’s multiple comparison test. ****p < 0.0001; ***p < 0.001; **p < 0.01; *p < 0.05; non-significant (ns).

Since both the U6 and U1 promoters used in this study are commonly employed for non-coding RNA expression ^35–37^, we investigated whether combining PTMs K1.2 and K6.5 with antisense sequences from asRNAs, all expressed under the U6 promoter in a single vector, would produce the same effect as when PTMs and asRNAs are expressed under the separate promoters. For the combined PTMs K1.2 and K6.5, we inserted the antisense sequence from the best-performing asRNAs upstream of the binding domain. We designed the combined PTMs with a 3-nucleotide spacer between the asRNA sequence and the binding domain to increase flexibility and allow both the binding domain and the antisense sequence to effectively bind to their respective target sites. However, flow cytometry results indicated that the combined PTMs K1.2 and K6.5 with asRNA in a single vector were less efficient than when PTMs and asRNAs were expressed under separate promoters and vectors. (Figure S2E and S2F).

Results obtained by flow cytometry were further confirmed by confocal microscopy where the strongest YFP signal was detected with the combination of PMT and asRNA (Figure S3B). Similarly, Western blot analysis of cell lysates upon immunostaining with anti-myc antibodies revealed bands corresponding to the full-length YFP (Figure 3A). Trans-splicing was also confirmed on a transcriptional level by semi-quantitative PCR. Upon co-transfection of HEK293T with the target intron reporter, corresponding best PTM candidates (K1.2, K5.2 or K6.5) and asRNA (BP4, SRP40, ASF21, SC35), RNA was isolated from the cell lysates and reverse transcribed to cDNA. Trans-splicing was confirmed through amplification of a segment by primers annealing to N-YFP and myc tag which are expected to co-localize in the trans-spliced reporter molecule (Figure 3B). The bands obtained upon the agarose gel electrophoresis corresponded to the expected size of the segment (Figure 3C) and Sanger sequencing of the segments extracted from the gel additionally confirmed that trans-splicing mediated joining of N-YFP and C-YFP segments took place at the RNA level (Figure 3E). No bands were observed in case of reporters lacking the target introns (Figure 3D). Next, a 265 bp segment of the N-YFP-C-YFP junction was amplified by qPCR to quantify the trans-splicing efficiency increase by asRNA (Figure 3B). The values were normalized to the expression of GAPDH and the trans-splicing with PMT alone. In accordance to the flow cytometry measurements, trans-splicing was strongly enhanced when PMT was combined with the asRNA. Specifically, a 10-fold increase in the YFP junction expression was noted with a combination of PTM K1.2 and BP4 compared to the K1.2 alone. PTM 6.5 and PTM K5.2 in combination with the respected asRNA led to a 5 to 10-fold increase in the YFP junction expression (Figure 3F, 3G and 3H). This increase was dependent on the presence of the target intron-bearing reporter (Figure 3F, 3G and 3H). A corresponding decrease in cis-splicing of the target intron reporter was also detected when we amplified a segment between N-YFP and the downstream CTNNB1 exon which are expected to remain co-localized in a cis-spliced target intron reporter (Figure 3I, 3J and 3K).

**Figure 3:**
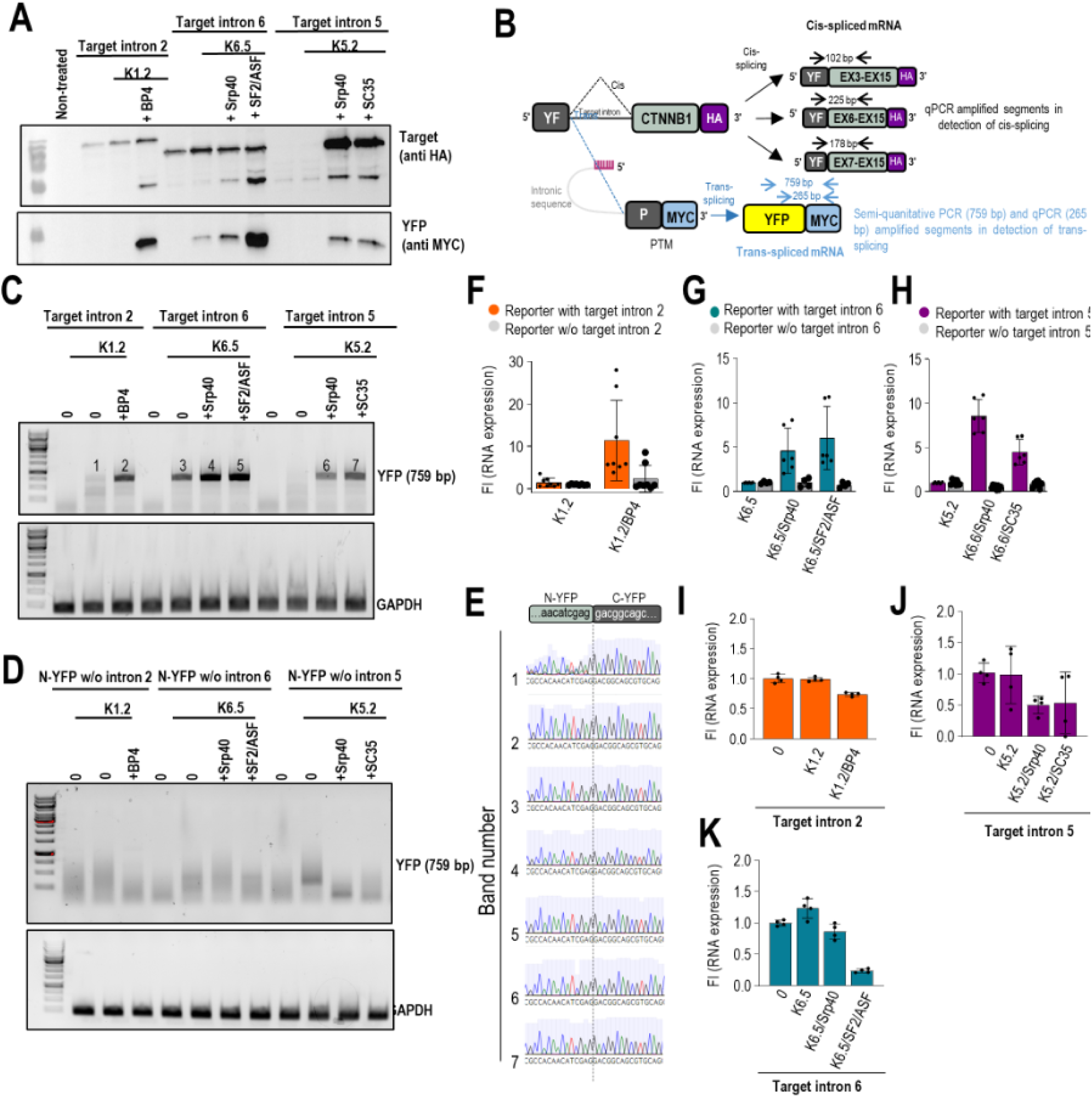
Detection of trans-splicing efficiency on mRNA and protein level. (A) Western blot depicting reconstituted YFP upon transRNA and asNA-facilitated trans-splicing of the reporters containing target introns 2, 6 and 5. Upon transfection cell lysates were subjected to SDS Page and WB with cis-spliced reporter detected via HA-tag and trans-spliced reporter detected via nYFP-myc-tag. Representative Western blot of two independent experiments is shown. (B) A schematic representation illustrating position and size of qPCR and semi-quantitative PCR amplified segments in detection of cis- and trans-splicing in (C-K). (C-E) Gel electrophoresis representing semi-quantitative PCR-amplified segments of nYFP-myc tag junction (759 bp) indicative of reporter trans-splicing. RNA was isolated from HEK293T cells and reverse-transcribed to cDNA. Bands numbered 1-7 (C) were isolated from the gel and submitted to Sanger sequencing with the obtained sequences corresponding to nYFP-myc tag junction (E). (F-H) qPCR-determined relative abundances of the RNA segment from the cYFP-myc tag junction (265 bp) indicative of trans-splicing. Fold increase is calculated compared to samples transfected with transRNA only. GAPDH was used as the reference and ΔΔCT method was used for quantification. (I-K) qPCR-determined relative abundances of the segment from the nYFP and cis-spliced exon junction indicative of cis-splicing. Fold increase is calculated compared to samples transfected with transRNA only. GAPDH was used as the reference and ΔΔCT method was used for quantification.

**Figure 4:**
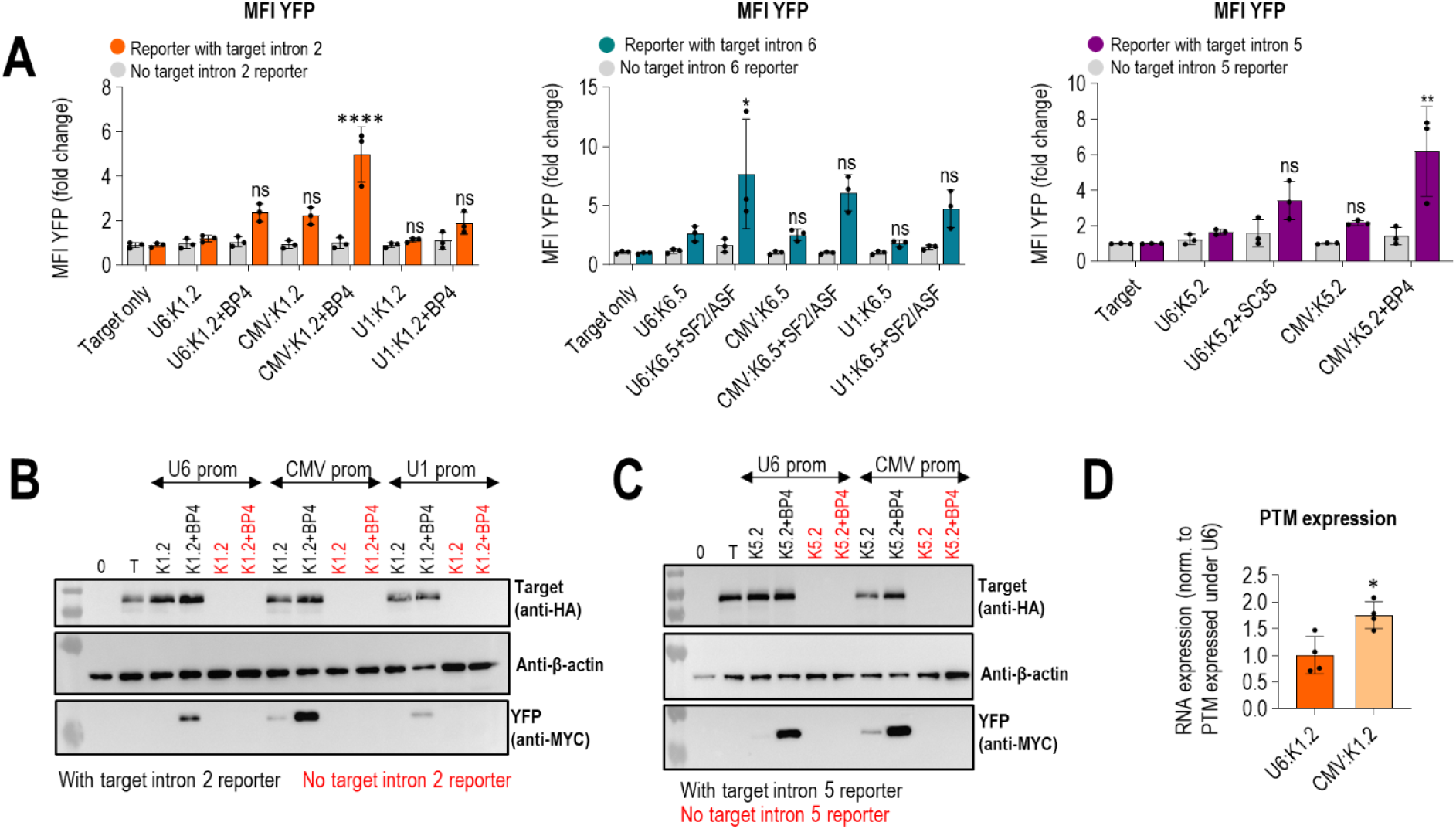
Expression of PTM K1.2 and K5.2 under CMV promoter improved trans-splicing efficiency. (A) Testing of PTM K1.2, K5.2 and K6.5 expressed under U6, CMV and U1 promoter for the trans-splicing efficiency using flow cytometry. Negative controls included target intron reporter transfected alone or PTM candidate transfected alone or with asRNA but without target intron reporter. Results are shown as YFP median fluorescence intensity (MFI) normalized to negative control – target intron reporter transfected alone. Data are presented as the mean value ± SD from at least three independent experiments. Comparison to PTM candidate co-transfected with target intron reporter was analyzed using one-way ANOVA with Dunnett’s multiple comparison test. ****p < 0.0001; **p ≤ 0.01 *p < 0.05; non significant (ns). (B, C) Trans-splicing efficiency of PTM K1.2 expressed under U6, CMV and U1 and PTM K5.2 expressed under U6 and CMV promoter was further verified with Western blot. Equal amounts (50 ug) of total proteins were loaded on SDS-PAGE and analyzed by immunoblotting using the corresponding anti-MYC, anti-HA and anti-β actin antibodies. 0 represents negative control – cells transfected with an empty plasmid vector (pcDNA3); T represents negative control – target intron reporter transfected alone; PTMs and asRNAs transfected with target intron reporter are marked in black; PTMs and asRNAs transfected without target intron reporter are marked in red. Data are representative of two independent experiments. (D) Comparison of PTM K1.2 expressed under U6 and CMV promoter by using qPCR with specific primers design to detect PTM expression. Expression of K1.2 from CMV was normalized to expression of K1.2 from U6 promoter. Comparison to K1.2 expressed under U6 promoter was analyzed using Student’s *t*-test (two populations) *p < 0.05; non significant (ns).

### Self-cleaving ribozymes at 5’ of the PMT significantly enhance 3’ trans-splicing

Next, we wanted to explore self-cleaving ribozymes as a tool to cleave off 5’ or 3’ ends of the PMT as the means of manipulating posttranscriptional processing such as polyadenylation and capping that importantly affect RNA trafficking from the nucleus, its stability and its translation ^38,39^. Ribozymes placed at the 3’ of the PMT have been shown to increase 5’ trans-splicing efficiency through cleavage of the polyadenylation tail ^30^. The polyadenylation tail of the PMTs designed for 5‘ trans-splicing, namely, is not trans-spliced into the target RNA and therefore only facilitates the undesirable nuclear export and translation of the PMT while being redundant for the translation of the trans-spliced RNA. We applied the analogous logic to our PMTs designed for 3‘ trans-splicing and assumed an equally redundant role of 5‘ positioned cap. In this, to our knowledge not yet utilized approach, we positioned various ribozymes on the 5‘ end of our best PMT candidates (K1.2, K6.5). We tested the effect of three self-cleaving ribozymes (Twister, genomic version of hepatitis D virus (HDV) and hammerhead (HH)), positioned either at 5’ or at 3’ of the two best PMT candidates K1.2 and K6.5 (Figure 5A).

**Figure 5:**
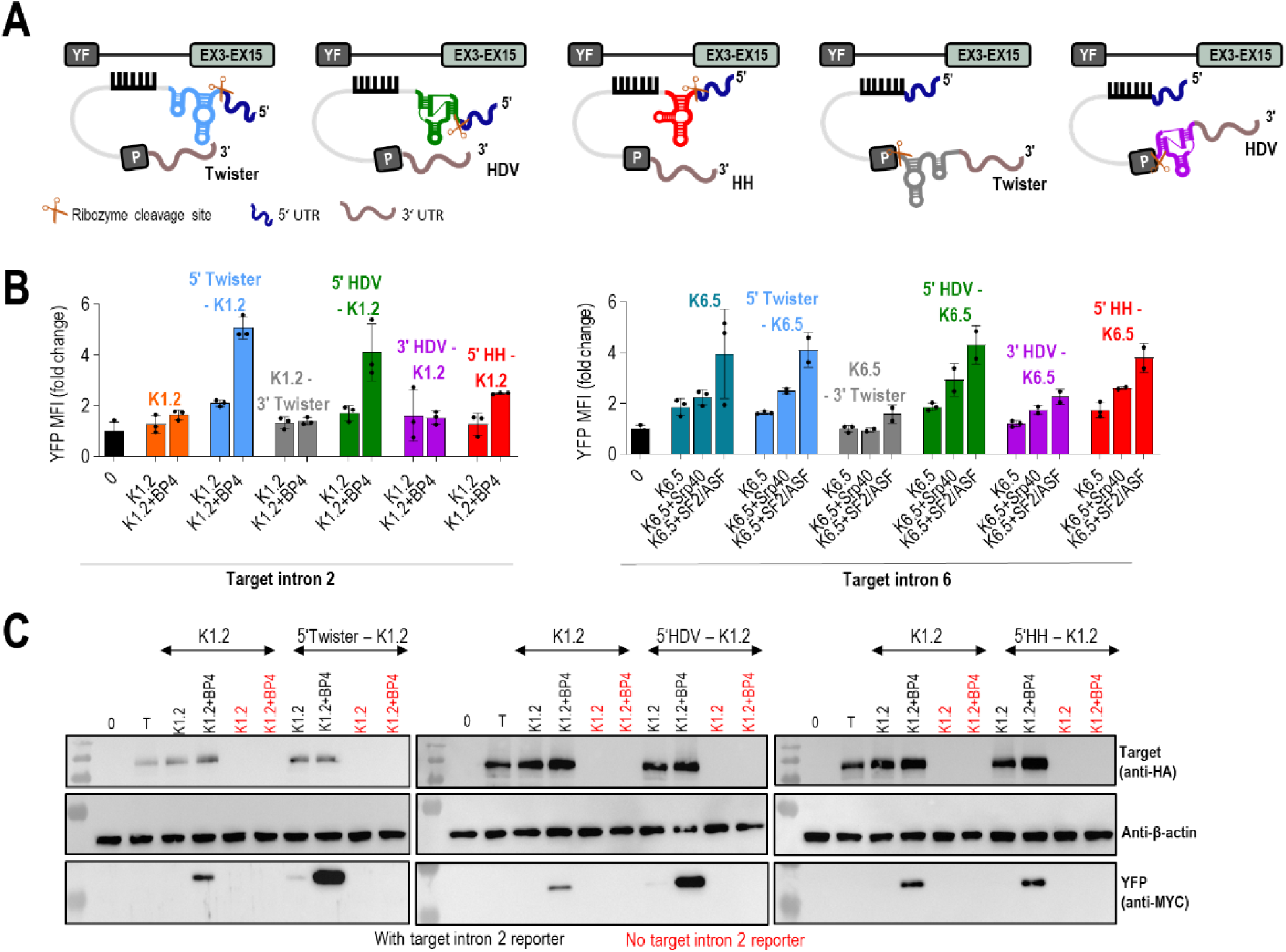
Addition of Twister and HDV ribozymes on 5’ end of PTM enhanced trans-splicing efficiency. (A) Schematic example of a PTM targeting intron 2 with added ribozymes on either 5’ or 3’ ends to remove capping or polyadenylation. (B) Determination of trans-splicing efficiency for the PTM K1.2 and PTM K6.5 by using flow cytometry. Negative control included the target intron reporter transfected alone. Quantitation by flow cytometry is presented as the median fluorescence intensity (MFI) of YFP normalized to the reporter-only control for the intron 2 and 6. Data are presented as the mean value ± SD from at least three independent experiments. (C) Detection of trans-splicing efficiency for the PTM K1.2 with ribozymes placed at 5’ end as restoration of full-length YFP by western blot analysis using the corresponding anti-myc, anti-HA and β-actin antibodies. 0 represents negative control – cells transfected with an empty plasmid vector (pcDNA3); T represents negative control – target intron reporter transfected alone; PTM k1.2 and asRNA BP4 transfected with target intron reporter are marked in black; PTM k1.2 and asRNA BP4 transfected without target intron reporter are marked in red. Data are representative of two independent experiments.

We observed a significant increase in the YFP signal with 5’ placement of the ribozymes on K1.2, which was further increased by the addition of asRNA BP4 (Figure 5B), These findings were corroborated by qPCR (Figure S4A) and Western blot analysis (Figure 5C). The ribozymes did not appear to additionally increase trans-splicing efficiency of K6.5 (Figure 5B). As a control, 3’ placement of ribozymes, expected to cleave the polyadenylation tail of the PMT, led to an increase in trans-splicing only at the RNA level (Figure S4A), likely due to the retention of the PMT inside the nucleus, but on a protein level we observed no additional increase in YFP signal with either of the target intron reporters (Figure 5B). This was expected due to the impaired translation of the trans-spliced RNA lacking the polyadenylation tail. Overall, these findings suggest that the truncation of the 5’ of PMT, which likely omits capping, is responsible for the increased efficiency.

To confirm that the improvement in trans-splicing efficiency observed after positioning the ribozyme to the 5’ end of the PTM molecule is indeed due to the ribozyme’s catalytic activity and not stabilization due to the formation of a terminal secondary RNA structure, we designed PTM K1.2 containing a 5’ HDV ribozyme with an inactivating mutation in its catalytic site. Flow cytometry analysis revealed that PTM K1.2 with the mutated HDV ribozyme exhibited reduced trans-splicing efficiency, particularly in the presence of the asRNA. The efficiency of PTM K1.2 containing the mutated ribozyme was comparable to that of PTM K1.2 without any prior optimization (Figure 6E and S4E). These results were further validated by quantitative PCR (qPCR), which corroborated the decrease in trans-splicing efficiency in the presence of the catalytically inactive ribozymes (Figure 6D). Additionally, the original unoptimized PMT K1.2 exhibited significant leakage to cytoplasm and translation into active β-catenin which was detectable by the TopFlash reporter assay (Figure 6I). β-catenin overexpression from the PMT is possible because it contains the majority of the coding CTNNB1 exons (3-15), however it is an undesired effect. Inclusion of the ribozyme significantly reduced translation of PMT, as evidenced by the TopFlash assay (Figure 6I), likely due to the increased retention of PMT within the nucleus and redirection into trans-splicing upon de-capping.

**Figure 6:**
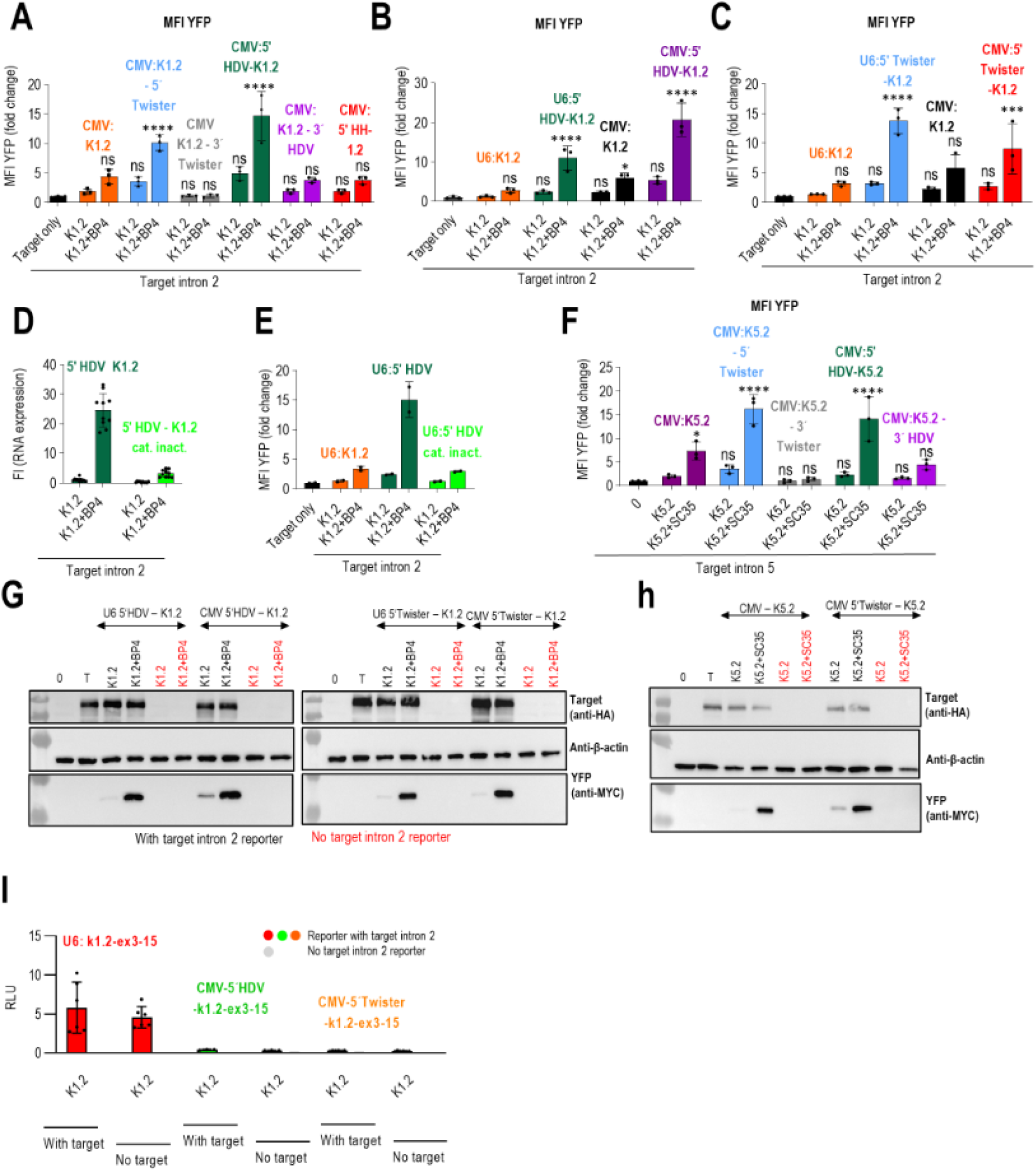
Combination of 5’ ribozymes and CMV expression of PTM K1.2 and K5.2 lead further to increased trans-splicing efficiency. (A) Testing PTM K1.2 expressed under CMV promoter with added HH, HDV and Twister ribozymes on either 5’ or 3’ end of PTM. Trans-splicing efficiency is detected as reconstitution of YFP detected with flow cytometry. Negative control (target only) included the target intron reporter transfected alone. Results are presented as the median fluorescence intensity (MFI) of YFP normalized to the reporter-only control. Data are presented as the mean value ± SD from at least three independent experiments. (B,C) Comparison of PTM expressed under CMV and U6 promoters with HDV (B) or Twister (C) added on 5’ end of PTM candidate. Trans-splicing was detected as YFP expression with flow cytometry. Results are shown as YFP MFI normalized to target intron reporter transfected alone. Data are presented as the mean value ± SD from at least three independent experiments. (D-E) Decrease in trans-splicing efficiency of PTM K1.2 bearing mutated HDV ribozyme on its 5’ end was measured via qPCR-determined relative abundance of nYFP-myc–tag junction (D) or by flow cytometric measurement of reconstituted YFP (E). (D) qPCR results are expressed as fold increase, calculated compared to samples transfected with transRNA only. GAPDH was used as the reference and ΔΔCT method was used for quantification. (E) Negative control (target only) for the flow cytometry included the target intron reporter transfected alone. Results are presented as the median fluorescence intensity (MFI) of YFP normalized to the reporter-only control. Data are presented as the mean value ± SD from at least three independent experiments. (F) Testing PTM K5.2 expressed under CMV promoter with added HDV and Twister ribozymes on either 5’ or 3’ end of PTM. Trans-splicing efficiency is detected as reconstitution of YFP detected with flow cytometry. Negative controls included the target intron reporter transfected. Results are presented as the median fluorescence intensity (MFI) of YFP normalized to the reporter-only control. Data are presented as the mean value ± SD from at least three independent experiments. (G, H) Detection of trans-splicing efficiency for the PTM K1.2 (G) or K5.2 (H) with HDV or Twister ribozyme placed at 5’ end as restoration of full-length YFP. Equal amounts (50 ug) of total proteins were loaded on SDS-PAGE and analyzed by immunoblotting using the corresponding anti-MYC, anti-HA and anti-β actin antibodies. 0 represents negative control – cells transfected with an empty plasmid vector (pcDNA3); T represents negative control – target intron reporter transfected alone. (G) PTM k1.2 and asRNA BP4 transfected with target intron reporter are marked in black; PTM k1.2 and asRNA BP4 transfected without target intron reporter are marked in red. (I) Relative luminescence upon co-transfection of TopFlash reporter plasmid with optimized K1.2 and/or target intron reporter with or without target intron. TopFlash reporter plasmid encodes luciferase expression under the β-catenin responsive promoter elements. To support dual luciferase assay, equal amounts of Renilla luciferase plasmid were added with each transfection mixture. (A-I) Data are representative of two independent experiments. Comparison to target intron reporter only (target only) was analyzed using one-way ANOVA with Dunnett’s multiple comparison test. ****p < 0.0001; ***p < 0.001; *p < 0.05; non significant (ns).

### Expression under the CMV promotor enhances trans-splicing at introns 2 and 5 targeting PMT

PMTs in the above-described experiments were expressed under the snRNA U6 promotor to support RNA stability and nuclear retention characteristics for non-coding RNAs^36^. To further improve trans-splicing efficiency, we compared the expression of best PTM candidates K1.2 and K6.5 under U1, U6 and CMV in the split YFP reporter system. Notably, K1.2 expressed under the CMV promotor exhibited better trans-splicing efficiency, as evidenced by both the percentage of YFP^+^ cells and mean fluorescence intensity (MFI), compared to the PTM expressed under U6 promotor (Figure 4A and S3A). We hypothesized that the increased trans-splicing efficiency observed with K1.2 expressed under the CMV promoter is since CMV is a stronger promoter than the U6 promoter is. We confirmed this by using qPCR with specific primers designed to detect PTM expression where we indeed detected higher expression of PTM K1.2 when expressed under CMV promoter (Figure 4D). A significant increase in YFP signal was also observed for the best-performing PTM, K5.2, expressed under the CMV promoter in combination with asRNA SC35 (Figure 4A and S3A). In contrast, CMV did not significantly improve trans-splicing by K6.5 PTM (Figure 4A and S3A). Improved trans-splicing by K1.2 and K5.2 PTMs expressed under CMV promoter were further validated by Western blot analysis (Figure 4B and 4C).

Given the strong trans-splicing enhancement of asRNA and PMT in comparison to PMT alone that was consistently observed (Figure 2D and 2E) we wondered, whether a single, hybrid molecule between asRNA and PMT could convey the same trans-splicing efficiency given the proximity between PMT BD and asRNA. For this purpose, all constituents of the PMT sequence except the BD (spacer, BP, ppt, 3’ ss, C-YFP and myc tag) were attached to the 5’ of the asRNA and expressed in its U1-U7 expression cassette. While such a construct supported trans-splicing, it was less efficient than the combination of PMT and asRNA from two separate constructs as well as the original PMT alone (Figure S3C) which indicates that a single chain may constrain binding to the two sites at the same time, which might be solved either by a longer linker of self-cleaving RNA.

After this optimization, we further examined the ability of PTM candidates to induce trans-splicing combining all enhancements that improved trans-splicing efficiency. Specifically, the best performing PTM candidate K1.2 was expressed under the CMV promoter, with self-cleaving ribozymes HH, HDV, and Twister added to either the 5’ or 3’. The optimized PTM K1.2 showed a further increase in trans-splicing efficiency on the target intron reporter when the HDV or Twister ribozymes were attached to the 5’ (Figure 6A and S4B). A similar effect was observed for the best-performing PTM K5.2, targeting intron 5. The addition of the Twister ribozyme to the 5’ end improved trans-splicing efficiency, as indicated by an increase in the percentage of YFP+ cells and MFI in flow cytometry (Figure 6F and S4D). Additionally, a combination of PTM K1.2 and PTM K5.2 with asRNAs also further enhanced trans-splicing efficiency, particularly as observed in MFI (Figure 6A, 6F and S4B, S4D), which indicates that each of these optimizations addresses different trans-splicing efficiency bottlenecks. Furthermore, results were validated with Western blot analysis (Figure 6G and 6H).

As we observed an improved trans-splicing in both cases when K1.2 was expressed under either the U6 or CMV promoter with HDV and Twister ribozymes placed at the 5’, we further compared these two K1.2 variants. The results showed a greater increase in trans-splicing efficiency with the 5’ HDV K1.2 expressed under the CMV promoter, while trans-splicing efficiency was similar for both promoters for the 5’ Twister K1.2. This conclusion is supported by results from flow cytometry (Figure 6B, 6C and S4C) and Western blot analysis (Figure 6G).

### Trans-splicing of endogenous CTNNB1

To assess ability of PTM candidates to induce RNA trans-splicing on endogenous CTNNB1 the 3’ C-YFP sequence of PTMs was replaced with the corresponding coding sequence of CTNNB1 depending on the intron insertion site with an added sequence for the myc tag at 3’. We initially transfected HEK293T cells with the most promising optimized PTM candidates for introns 2, 5 and 6 and successful trans-splicing resulted in the fusion of endogenous CTNNB1 pre-mRNA upstream from the target intron with the wild-type coding sequence of CTNNB1-Myc derived from the PTM (Figure 7A and S5A). Forty-eight hours after transfection total RNA was isolated and reverse transcribed. Semi-quantitative PCR with specific forward primer for endogenous CTNNB1 and a specific reverse primer for MycTag produced trans-spliced PCR fragments of the predicted size 2435 bp for the PTM CMV:5’HDV:K1.2, 1575 bp for the PTM U6:K6.5 and 1881 bp for the CMV:5’Twister:K5.2. These primers do not generate PCR products from a cis-spliced target. The successful specific trans-splicing was confirmed for intron 2. Furthermore, trans-splicing was detected when asRNAs were added in combination with PTM candidate (Figure 7B). The sequence analysis of trans-spliced fragments further validated the occurrence of trans-splicing (Figure 7C). In case of PTM K5.2 and K6.5 we were not able to detect specific on-target trans-splicing on mRNA level (Figure S5B and S5C).

**Figure 7:**
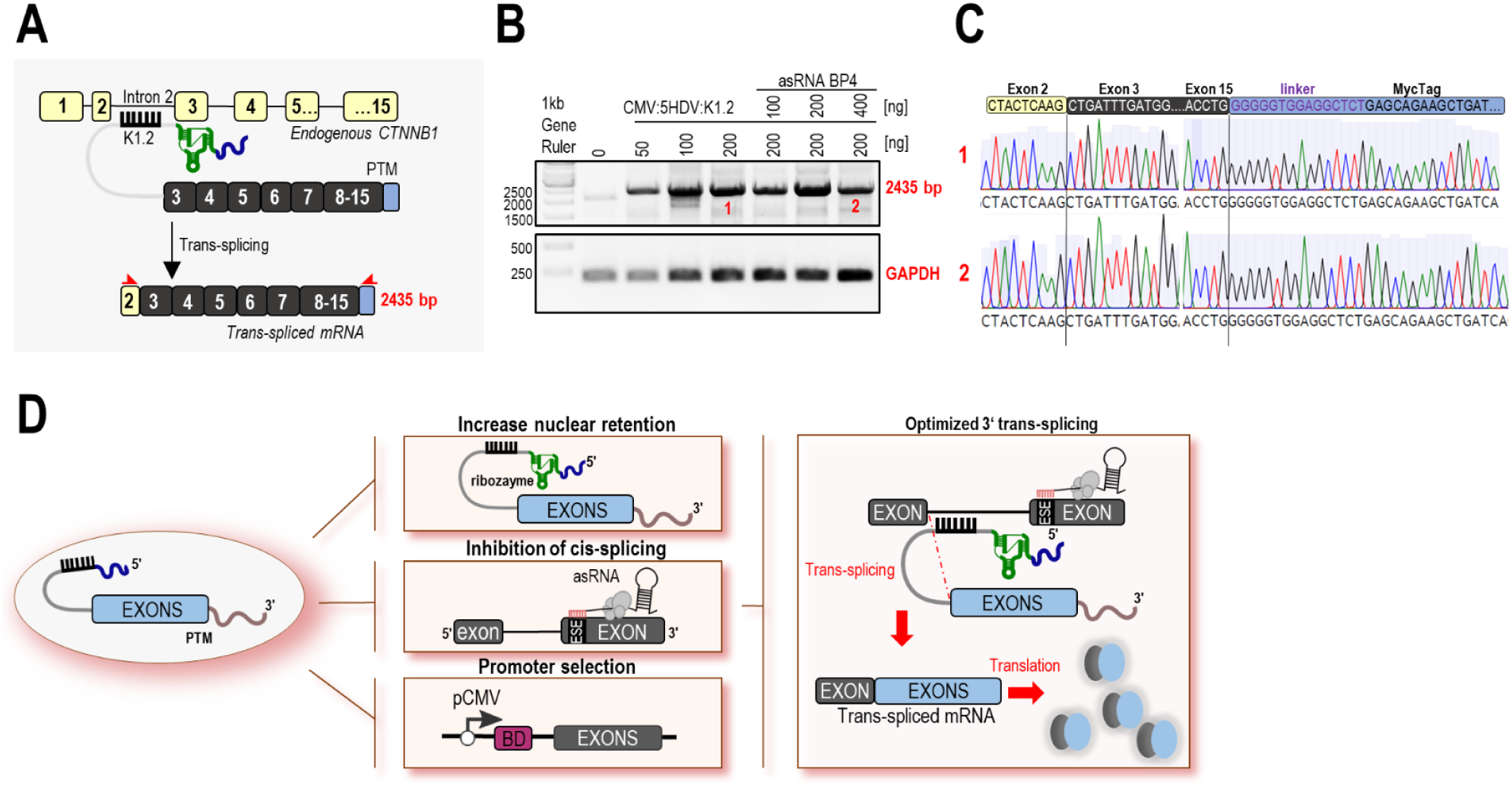
Endogenous detection of trans-splicing efficiency in HEK293T cells. (A) Schematic example of best performing PTM K1.2 and PTM K5.2 with coding CTNNB1 region targeting endogenous CTNNB1 target introns 2 and 5. Result of trans-splicing is trans-spliced CTNNB1 mRNA that contains endogenous CTNNB1 region upstream from target intron (yellow exons) and exogenous CTNNB1 region with myc tag from PTM candidate. To detect trans-spliced mRNA by using PCR, specific primers (asterisks marked in red) were design in order to detect exon 2 and myc tag for the PTM K1.2 and exon 5 and myc tag for the K5.2. Size of bands that corresponds to trans-spliced mRNA is 2435 bp for the K1.2 and 1896 bp for the PTM K5.2. (B) Correctly trans-spliced CTNNB1 mRNA was detected 48h after transfection by using PCR. Size of trans-spliced CTNNB1 product is 2435 bp for the K1.2. HEK293T cell transfecting with empty vector (pcDNA3) were used as negative control. (C) Sanger sequencing of PRC product further confirmed accuracy of the trans-splicing in HEK293T cells. Bands labeled 1 and 2 were excised from the gel and subsequently sent for Sanger sequencing. Data are representative of two independent experiments. (D) Schematic representation of the best optimizations performed in this study to increase trans-splicing efficiency. The novel implementation of 5’ ribozymes in PTMs expressed under the CMV promoter, in combination with asRNA to block cis-splicing, improved trans-splicing efficiency.

## Discussion

Gene therapy strategies like gene replacement therapy and regulation of gene expression with siRNA and antisense oligonucleotides (ASO) and CRISPR-Cas genome editing techniques are increasingly entering clinics. There is a pressing need to expand the repertoire of advanced therapies with strategies that can provide safety due to the reliance on endogenous regulation. The RNA transcript-targeting technique of spliceosome-mediated pre-mRNA trans-splicing (SMaRT) holds the potential to circumvent many of the challenges of other methods particularly where excessive expression could be toxic. While the SMaRT-based therapy for ABCA4-related retinopathy is currently being evaluated in a clinical trial^40^ its implementation for other targets and improvement of the efficiency of SMaRT are very much needed. Here, we explored the potential of SMaRT strategy with novel modifications aiming for the therapeutic strategy for the CTNNB1 syndrome. Upregulation of the endogenous CTNNB1 gene expression up to 2-fold from the remaining functional gene copy should be sufficient to rescue its function while the gene replacement therapy typically yields much higher upregulation. In some genes, including the neurodevelopmentally important CTNNB1, FOXG1 and EHMT1, this is problematic due to their sensitivity to both under- as well as overexpression, which requires their gene expression to be strictly maintained within a narrow window. As the SMaRT strategy does not interfere with the gene regulation but rather corrects the effects of the mutation on a transcriptional level, dosage-sensitive genes such as CTNNB1 are ideal candidates for its implementation.

The selection of suitable target introns within the CTNNB1 transcript was guided by 3’ splice site strength predictions, which indicated that introns 2, 3, 5, and 6 had lower 3’ splice site strength then the PTM, making them ideal candidates for trans-splicing. We employed a tiling approach to design PTM with varying binding domains (BDs) across the target introns, which allowed us to assess which BDs provided the highest trans-splicing efficiency. This approach revealed the three most promising PTM candidates for introns 2, 5 and 6. It has been reported that the position of PTM binding in the target intron correlates with the trans-splicing efficiency. In accordance with other 3’ trans-splicing adopting studies^41,42^, binding domains that target 5’ part of the target intron showed better trans-splicing efficiency as also exhibited by our best PTM candidates, K1.2 and K6.5. The proximity of the PTM candidate to the 5’ splice site likely allows the spliceosome machinery to recognize the 3’ splice site of the PTM candidate before the 3’ splice site of the target intron, which consequently increases the likelihood of inducing trans-splicing. The addition of short antisense RNAs (asRNAs) designed to target 3’ splice site and exon splicing enhancers (ESE) significantly boosted trans-splicing efficiency for all tested introns. This enhancement was most notable for intron 5, where the asRNA SC35 increased YFP^+^ cells from 16% to 55%. These findings are in line with previous studies which utilized short synthetic antisense oligonucleotides to inhibit competing cis-splicing, thereby promoting the desired trans-splicing event^22^. However, combining multiple asRNAs did not result in further improvement, indicating that a single, well-designed asRNA is sufficient to enhance PTM-mediated trans-splicing.

The choice of promoter also played an important role in determining the trans-splicing efficiency. Expressing PTM K1.2 under the CMV promoter significantly enhanced trans-splicing compared to the initial choice of U6 promoter, while K6.5 showed no significant difference between promoters. CMV promoter is utilized by PolII and results in a capped RNA. We further explored the use of self-cleaving ribozymes at the 5’ and 3’ ends of the PTM molecules. Placement of an active self-cleaving ribozyme at the 5’ end of the PTM K1.2 and PTM 5.2 significantly enhanced trans-splicing efficiency on both RNA and protein level and significantly reduced the leakage of PTM 1.2 into translation, likely due to the removal of the 5’ cap. Ribozymes have thus far been utilized to cleave the polyadenylation tail off the 5’ trans-splicing PTMs to facilitate their retention inside the nucleus ^30^. Our findings are the first to suggest that 3’ trans-splicing can be analogously enhanced by cleaving the redundant 5’ cap of PMT. Transcripts from both polymerase II and III driving transcription from CMV and U6 promotors, respectively, were reported to possess a 5‘ cap (7-methyl guanine^39^ and gamma monomethyl^43^). Accordingly, in our experiments cleaving off the 5‘ cap from the PMTs expressed under both promotors significantly enhanced 3‘ trans-splicing. Finally, using combined optimizations we successfully demonstrated that PTM targeting intron 2 induce robust on-target trans-splicing of the endogenous CTNNB1 transcripts in HEK293T cells. While best results were obtained from a combination of PTM with ribozyme and asRNA in separate constructs, future efforts may be able to provide a single construct strategy that could be more robust for therapeutic applications. The experiments focused on CTNNB1, although the same design rules are likely to be applicable for other therapeutic targets.

In conclusion, this study presents an exploration of strategies to optimize PTM-mediated trans-splicing for the CTNNB1 gene. Our findings underscore the importance of combining the selection of the binding domain closer to the 5‘ end of the target intron, the addition of asRNA to mask cis-splicing motifs, the use of CMV promotor, and cleavage of 5‘ of the 3‘ trans-splicing PMT in achieving high efficiency of trans-splicing (Figure 7D). These results provide a solid foundation for future applications of trans-splicing for the CTNNB1 syndrome, including gene editing and the treatment of genetic disorders.

## Materials and methods

### Plasmid construction

Split YFP target minigenes were cloned into the pcDNA3.1D vector backbone. PTM are cloned into the pgRNA or pcDNA3.1D vector backbone. The plasmid encoded with CTNNB1 was provided by OriGene Technologies GmbH, Herford Germany. Target CTNNB1 intronic sequences were ordered as eblock (IDT, Inc., Coralville, Iowa, USA). PCR amplification was performed using RepliQa HiFi ToughMix® (Quantabio, Beverly, MA, USA). All plasmids were constructed using the Gibson assembly method and are provided in Table S5.

### Construction of asRNA library

ESEfinder and SVM-BPfinder prediction tools were used to predict exonic splicing enhancer (ESE) sequences and branch point (BP). Sequence for the cassette with U1 promotor and modified U7 snRNA was taken from the literature^32^ and ordered as eblock (IDT, Inc., Coralville, Iowa, USA) and cloned into pgRNA vector (Addgene plasmid #44248) using Gibson assembly method. Designed asRNA sequences were ordered as primers and added to U1U7 cassette using Gibson assembly. Detailed sequences of all asRNA constructs are listed in **Table S5**.

### Cell culture

HEK293T cells (American Type Culture Collection) were cultured in DMEM medium (Thermo Fisher Scientific) supplemented with 10% v/v FBS (Thermo Fisher Scientific). Cell line was maintained in a humidified incubator at 37°C with 5% CO2. Cell lines were obtained from the ATCC culture collection.

### Transient transfection

HEK293T cells were washed with phosphate-buffered saline (PBS) and detached from the surface using Trypsin-EDTA solution (Sigma-Aldrich; T3924). The cell concentration was measured using Countess™ Cell Counting Chamber Slides or EVE™ Cell Counting Slides kits with Trypan blue as an indicator of live cells and measured on the Countess™ 3 Automated Cell Counter (Invitrogen™). For RNA extraction, immunoblotting and flow cytometry, 1×10^5^ cells per well were seeded in 24-well TPP plates. At 30%–50% confluence, HEK293T cells were transfected with a mixture of DNA and PEI (6 µl per 500 ng of DNA, stock concentration 0.324 mg/ml, pH 7.5). The PEI stock concentration was diluted in 150 mM NaCl and mixed at a 1:1 ratio with the appropriate DNA, also diluted in 150mM NaCl. This was incubated at room temperature for 15 min and added to the cell media in plates. The amounts of transfected plasmids are listed in **Table S4**. For cytometry experiments, plasmid expressing iRFP was added to transfection mixes as a transfection control.

### Flow cytometry

The transfected HEK293T cells were maintained at 37 °C in a 5% CO2 environment. At 48 h post-transfection, the cells were prepared in PBS for the analysis on the Cytek™ Aurora Flow Cytometry System (Cytek® Biosciences, Fremont, California, US). For flow cytometry controls and unmixing, we used non-transfected HEK293T cells as negative control and constitutive expression vectors encoding YFP, tagBFP, and iRFP fluorescent proteins as single stain controls. Cells were first gated for singlets, then the expression of iRFP as transfection control and then for the expression of YFP. Gating strategy is described in **Figure S6.**

### RNA isolation and cDNA synthesis

At 48 hours, RNA was extracted from transfected HEK293T using the High Pure RNA Isolation Kit (Roche, Basel, Switzerland) according to the manufacturer’s protocol. Reverse transcription was performed on 1µg of total RNA using a high-capacity complementary DNA (cDNA) reverse transcription kit (Applied Biosystems, Waltham, Massachusetts, USA) with a mixture of random oligonucleotides, according to the manufacturer’s instructions.

### Semi-quantitative PCR

To detect trans-splicing efficiency on mRNA level, semi-quantitative PCR amplification was performed on the cDNA using RepliQa HiFi ToughMix® (Quantabio, Beverly, MA, USA). Primers we designed to amplify the 759 bp long segment between N-YFP and myc tag that is assembled only in the trans-spliced PMT. PCR bands were analyzed via gel electrophoresis on the 1 % agarose gel (Zellbio, Lonsee, Baden-Württemberg, Germany), extracted and purified by peqGOLD gel extraction kit (VWR Peqlab, Darmstadt, Germany) and analyzed by Sanger sequencing.

### Western blot analysis

The cells were washed with PBS and lysed in 100 µl of 1× Passive lysis buffer (Promega, Madison, Wisconsin, US). Total protein concentration in the supernatant was measured by the bicinchoninic acid (BCA) method. Samples were denatured by 5-minute incubation at 95 °C in SDS with reducing agent. We then loaded 30-50 µg of total protein per sample onto the SDS–PAGE gel (Bio-Rad, Hercules, California, US) with PageRuler™ Plus Prestained Protein Ladder (Thermo Scientific™, Waltham, MA, US) as size standard. SDS-PAGE was run under denaturizing conditions at 200 V for 60 min. Proteins were transferred to a Hybond ECL nitrocellulose membrane (GE Healthcare, Chicago, Illinois, US). Myc-tagged trans-spliced YFP was specifically detected with primary antibodies Mouse anti-Myc (Cell Signaling Technology, Cat. No. 2276S) at 1:1000 and secondary antibodies Goat anti-mouse-HRP (Jackson ImmunoResearch, Cat.No. 115-035-003) at 1:2000 ratios. HA-tagged target reporter was specifically detected with primary antibodies Rabbit anti-HA-tag (Sigma-Aldrich, Cat.No. H6908) at 1:1000 and secondary antibodies Goat anti-rabbit-HRP (Thermo Fischer Scientific, Cat.No. 65-6120) at 1:2000 ratios. For loading control, we detected β-actin using antibodies Mouse anti-β-actin (Cell Signalling Technology, Cat.No. 3700) at 1:1000 and secondary antibodies Goat anti-mouse-HRP at 1:3000 ratios or α/β tubulin protein using primary antibodies Rabbit Alpha/Beta Tubulin (Cell Singlaing Tchnology, Cat.No. 2148) at 1:1000 ratios and secondary antibodies Goat anti-rabbit-HRP at 1:2000 ratios. Detection of HRP was achieved by incubation of the membrane with SuperSignal™ West Pico or Femto Maximum Sensitivity Substrate (Thermo Scientific™, Waltham, MA, US). The immunoblots were visualized on G-box membrane was imaged with G:Box Chemi XT 4 Chemiluminescence and Fluorescence Imaging System (Syngene, Bangalore, Karnataka, India).

### Confocal microscopy

Co-transfection of PMT and asRNA was performed in 8-well IBIDI plates for 48h. Microscopic images of cells were taken by a Leica TCS SP5 inverted laser-scanning microscope on a Leica DMI 6000 CS module equipped with an HCX Plan-Apochromat lambda blue 63× objective, numerical aperture 1.4 (Leica Microsystems, Wetzlar, Germany). A 488 nm laser line of 100 mW argon laser with 10 % laser power was used for the detection of YFP, where emitted light was detected between 500 and 600 nm. For acquisition and image processing we used Leica LAS AF program (Leica Microsystems, Wetzlar, Germany).

### Quantitative PCR (qPCR)

Co-transfection of PMT and asRNA into HEK293T was performed in 24-well format for 48h. Upon RNA isolation and reverse transcription to cDNA we determined the efficiency of trans- and cis-splicing using qPCR. To detect trans-splicing efficiency, primers were designed to amplify the 265 bp long segment between N-YFP and myc tag that is assembled only in the trans-spliced PMT. To detect cis-splicing, primers were designed to amplify the segment between N-YFP and the exon of the coding region that follows each target intron (Fig. 3b). A white 96-well LightCycler 480 Multiwell Plate was used for qPCR reaction with each reaction containing 1X SYBR Green I Master Mix (Roche, Basel, Switzerland), 20 ng of cDNA and 0.5 µM of forward and reverse primer and was run in LightCycler 480 instrument (Roche, Basel, Switzerland). As a control of the endogenous gene expression we amplified a short segment of GAPDH. All measurements were performed in duplicates or triplicates. Relative change in RNA expression was calculated using the 2-ΔΔCq method.

### Statistics

Results were analysed in Excel and presented in GraphPad Prism (GraphPad, Boston, US). Statistical significance was determined by the one-way ANOVA with the Dunnett ’s multiple comparisons test and Student’s t-test test. Results with p-value < 0.05 were deemed statistically significant.

## Data availability statement

All data supporting the findings of this study are available within the article or Supplemental Information. Additional data are available from the corresponding author upon reasonable request.

## Acknowledgments

This work was funded by the Slovenian Research Agency Z1-3193 (to PS-L), J7-4537 and P4-0176 and CTNNB1 Foundation.

## Author contributions

MM conceptualized the study, designed and performed the experiments and analysed the data, PS-L conceptualized the study, designed and performed the experiments, analysed the data as well as obtained funding. MMe performed the experiments. RJ conceptualized and supervised the study.

## Abbreviations

asRNA: antisense RNA
BD: Binding domain
BP: branch point
CTNNB1: catenin beta 1
ESE: Exonic splicing enhancer
MFI: Mean fluorescence intensity
mRNA: Messenger RNA
qPCR: quantitative PCR
PMT: pre-RNA trans-splicing molecule
pre-mRNA: precursor messenger RNA
SMaRT: Spliceosome-mediated RNA trans-splicing
YFP: Yellow fluorescent protein

